# Network States in the Basolateral Amygdala Predicts Voluntary Alcohol Consumption

**DOI:** 10.1101/2023.06.21.545962

**Authors:** Alyssa DiLeo, Pantelis Antonodiou, Katrina Blandino, Eli Conlin, Laverne Melón, Jamie L. Maguire

## Abstract

Although most adults in the United States will drink alcohol in their life, only about 6% will go on to develop an alcohol use disorder (AUD). While a great deal of work has furthered our understanding of the cycle of addiction, it remains unclear why certain people transition to disordered drinking. Altered activity in regions implicated in AUDs, like the basolateral amygdala (BLA), has been suggested to play a role in the pathophysiology of AUDs, but how these networks contribute to alcohol misuse remains unclear. Our recent work demonstrated that alcohol can modulate BLA network states and that GABAergic parvalbumin (PV) interneurons are crucial modulators of network activity in the BLA. Further, our lab has demonstrated that δ subunit-containing GABA_A_ receptors, which are modulated by alcohol, are highly expressed on PV interneurons in the BLA. These receptors on PV interneurons have also been shown to influence alcohol intake in a voluntary binge drinking paradigm and anxiety-like behavior in withdrawal. Therefore, we hypothesized that alcohol may impact BLA network states via δ subunit-containing GABA_A_ receptors on PV interneurons to impact the extent of alcohol use. To test this hypothesis, we measured the impact of acute alcohol exposure on oscillatory states in the basolateral amygdala and then assessed the relationship to the extent of voluntary ethanol consumption in the Intermittent Access, Drinking-in-the-Dark-Multiple Scheduled Access, and Chronic Intermittent Ethanol exposure paradigms. Remarkably, we demonstrate that the average alcohol intake negatively correlates with δ subunit-containing GABA_A_ receptor expression on PV interneurons and gamma power in the BLA after the first exposure to alcohol. These data implicate δ subunit-containing GABA_A_ receptor expression on PV interneurons in the BLA in voluntary alcohol intake and suggest that BLA network states may serve as a useful biomarker for those at risk for alcohol misuse.

**Significance Statement:** Oscillatory states in the BLA have been demonstrated to drive behavioral states involved in emotional processing, including negative valence processing. Given that negative emotional states/hyperkatifeia contribute to the cycle of AUDs, our previous work demonstrating the ability of alcohol to modulate BLA network states and thereby behavioral states suggests that this mechanism may influence alcohol intake. Here we demonstrate a relationship between the ability of alcohol to modulate oscillations in the BLA and future alcohol intake such that the extent to which alcohol influences BLA network states predict the extent of future voluntary alcohol intake. These findings suggest that individual variability in the sensitivity of the BLA network to alcohol influences voluntary alcohol consumption.

## Introduction

Approximately 85% of adults in the United States will drink alcohol in their lifetime and about 6% will develop an alcohol use disorder (AUD) (SAMHSA, 2018). Despite the large societal and health impacts, it is still unclear why certain people transition to disordered drinking (Koob & Volkow, 2009). The BLA is recognized as an integrative hub for processing emotional and rewarding stimuli, playing a role in fear, anxiety, social behaviors, reward, and motivation (Janak & Tye, 2015; Schoenbaum, Chiba, & Gallagher, 1999; Tye et al., 2011) and has, and has been implicated in the pathophysiology of AUDs and anxiety disorders (Agoglia and Herman, 2018; Silberman et al., 2009; Tye et al., 2011). The BLA is made up of mostly excitatory, glutamatergic principal cells and a heterogenous population of local GABAergic interneurons (Capogna, 2014; Muller, Mascagni, & McDonald, 2006). Increased activity in the BLA has been established as a hallmark feature of mood disorders, especially in those that are highly comorbid with AUDs, such as anxiety (Etkin & Wager, 2007; Rau, Chappell, Butler, Ariwodola, & Weiner, 2015; Zhou, Wang, & Zhu, 2010). Additionally, the BLA is important for reward salience, motivation, and drug associated cues, which play a role in relapse (Chaudhri, Woods, Sahuque, Gill, & Janak, 2013; Sciascia et al., 2015; Wassum & Izquierdo, 2015). Clearly, BLA activity lies at the crossroads of alcohol use disorders and mood disorders, highlighting our interest in the area.

Disturbed network communication in neuropsychiatric disorders is emerging as a theory for behavioral symptom manifestation (Herrmann & Demiralp, 2005; Uhlhaas & Singer, 2012; Voytek & Knight, 2015). Further, EEG signals are being pursued as a potential predictive biomarker to assist in diagnosis and treatment. For example, gamma oscillations are an important biomarker for schizophrenia and major depression (Fitzgerald & Watson, 2018; McNally & McCarley, 2016) and EEG recordings are being used to predict responses to drug treatment (Javitt, Spencer, Thaker, Winterer, & Hajos, 2008; Wu et al., 2020; Zhdanov et al., 2020). Relevant to alcohol use, EEG studies in human drinkers have found distinct differences in electrical activity between non-drinkers and drinkers (Courtney & Polich, 2010; López-Caneda et al., 2017; Rangaswamy et al., 2003). Further, alcohol also affects resting-state network connectivity attributed to inhibitory neurotransmission in humans (Lithari, Klados, Pappas, Albani, & Kapoukranidou, 2012). Preclinically, recordings of neuronal activity have been used to predict alcohol consumption (Hernandez & Moorman, 2020; Siciliano et al., 2019). However, how network states contribute to AUDs remains unclear, which may further our understanding of the underlying pathophysiology of disease as well as identify potential biomarkers for risk.

Network oscillations are a means of communication within the brain and are emerging as critical mediators of behavior, highlighting the importance of understanding the neural mechanisms of oscillatory communication (Buzsáki & Draguhn, 2004; Fitzgerald & Watson, 2018; Gonzalez-Burgos, Cho, & Lewis, 2015). Previous work from our lab and others have demonstrated that specific oscillation frequencies in the BLA orchestrate fear and safety states and are impacted by chronic stress and pharmacological treatments (Antonoudiou et al., 2022; Davis et al., 2017; Likhtik et al., 2013; Ozawa et al., 2020; Stujenske, Likhtik, Topiwala, & Gordon, 2014). Given that the BLA plays a critical role in mediating fear and anxiety (Janak & Tye, 2015), as well as alcohol consumption (Moaddab, Mangone, Ray, & Mcdannald, 2017; Sciascia, Reese, Janak, & Chaudhri, 2015; Varodayan et al., 2017), we hypothesize that similar network activity may influence drinking behavior. Thus, understanding how alcohol affects oscillations in the BLA and influences alcohol consumption may identify potential biomarkers, as well as novel treatments for AUDs.

The generation of oscillations is thought to involve GABAergic interneurons, such as PV expressing interneurons (Bartos, Vida, & Jonas, 2007; Fuchs et al., 2007; Sohal, Zhang, Yizhar, & Deisseroth, 2009). PV interneurons are fast spiking cells that make inhibitory contact on somas of principal cells, allowing them to exert powerful control over their activity (Bocchio et al., 2016; Muller et al., 2006). Unlike other interneurons in the brain, the fast intrinsic signaling properties of PV cells lend them to generating gamma oscillations between 30-100 Hz (Sohal et al., 2009), which are thought to underlie network communication in cognition, memory, and attentional tasks (Buzsáki & Wang, 2012; Colgin et al., 2009; Fries, 2005; Gruber, Tsivilis, & Muller, 2003). In fact, shifts in oscillation frequency in the BLA, between pro-fear (3-6 Hz) or pro-safety (6-12 Hz) states, are mediated by PV interneurons and impact behavioral expression of fear and anxiety and the behavioral consequences of chronic stress (Antonoudiou, 2022; Davis et al., 2017; Likhtik et al., 2013; Stujenske et al., 2014). In addition, we have shown that PV interneurons mediate the ability of norepinephrine to modulate gamma oscillations in the BLA which is associated with enhanced fear learning (Fu and Teboul, 2022). Our laboratory has demonstrated a critical role for PV interneurons in contributing to oscillations within the BLA (Davis et al., 2017; Antonoudiou, 2022). Further, we have shown that PV interneurons in the BLA express a high density of δ-GABA_A_ receptors (DiLeo, 2022), which confer sensitivity to alcohol (Glykys et al., 2007; Wallner, Hanchar, & Olsen, 2003; Hanchar et al., 2006; Santhakumar, Wallner, & Otis, 2007; Sundstrom-Poromaa et al., 2002) and have been shown to modulate hippocampal oscillations (Ferando & Mody, 2013, 2015; Mann & Mody, 2010; Pavlov et al., 2014). We propose that δ-GABA_A_ receptors expressed on PV interneurons in the BLA may modulate oscillations in the BLA and play a role in network synchronization (Pavlov et al., 2014) related to behavioral states, including alcohol consumption.

Our previous work demonstrated that alcohol is capable of modulating BLA network states in response to acute alcohol exposure which involves a role for δ-GABAA receptors (DiLeo, 2022). Here we demonstrate that the response of the BLA network states to the first exposure to alcohol predicts their future level of voluntary alcohol intake. These data suggest that there are inherent differences in the BLA network which underlies the extent of alcohol use. Our findings suggest that differences in the expression of δ-GABAA receptors in the BLA correlates with alcohol intake and may impact the ability of alcohol to impact BLA network states and influence alcohol consumption.

## Methods

### Animals

Adult male and female C57BL/6J mice, aged 8-12 weeks old, were purchased from The Jackson Laboratory (stock #000664) and group-housed in temperature and humidity-controlled housing rooms on a 12-hour light-dark cycle (lights on at 7 AM) with ad libitum food and water. Animals were handled according to protocols and procedures approved by the Institutional Animal Care and Use Committee (IACUC). Female mice were maintained in an acyclic state without exposure to males. Mice were singly housed and habituated to new cages for 24 hours before the start of experiments.

### Stereotaxic Surgery

All mice undergoing surgery were anesthetized with ketamine/xylazine (90 mg/kg and 5-10 mg/kg, respectively, i.p.) and treated with sustained-release buprenorphine (0.5-1.0 mg/kg, subcutaneously, s.c.). A lengthwise incision was made to expose the skull, and a unilateral craniotomy was performed to lower a depth electrode (PFA-coated stainless-steel wire, A-M systems) into the BLA (AP -1.50 mm, ML 3.30 mm, DV -5 mm), affixed to a head mount (Pinnacle #8201) with stainless steel screws as ground, reference, and frontal cortex electroencephalographic (EEG) (AP +0.75 mm, ML ± 0.3 mm, DV -2.1 mm) electrodes. Electromyographic (EMG) wires were positioned in the neck muscles.

### LFP Recordings

Local field potential (LFP) recordings were performed in male and female C57BL/6J mice after a week of recovery from implant surgery. LFP recordings were acquired using Lab Chart software (AD Instruments), collected at 4 kHz, and amplified 100X. Spectral analysis was performed in MATLAB (Antonoudiou et al., 2022) using MatWAND (https://github.com/pantelisantonoudiou/MatWAND), which utilizes the fast Fourier transform similar to previous reports (Freeman et al., 2000; Frigo & Johnson, 1998; Kruse & Eckhorn, 1996; H. C. Pape & Driesang, 1998). Briefly, recordings were divided into 5-second overlapping segments, and the power spectral density for a range of frequencies was obtained (Oppenheim et al., 2010). LFP power was quantified, analyzed, and graphed as power area (PA). Further, the PA change or difference (PA Diff) was calculated to measure how the LFP PA changed across treatment periods.

### Acute Alcohol Exposure

Before starting the experimental paradigm, mice were habituated to new cages with ad libitum food and water for 24 hours. All injections were performed 2-3 hours into the light cycle (on at 7 AM) at the same time each day across all cohorts. The acute exposure consisted of a baseline period followed by a saline injection (0.9% NaCl i.p.) and a subsequent 1 g/kg ethanol (EtOH) injection (20% *v/v* i.p.) as previously described (DiLeo, 2022). Each recording period was of minimum of 60 minutes.

### Intermittent Access

Adult male and female C57BL/6J mice underwent an intermittent access two bottle choice (IA2BC) paradigm for three weeks. On Mondays, Wednesdays, and Fridays, mice received a bottle of alcohol and water for 24 hours in their home cage three hours into their light cycle and removed at the same time the following day. Bottles were measured to the nearest tenth of a gram each day. To control for evaporation and spillage, drip bottles were designated in empty cages, and the average loss per week was subtracted from individual measurements. During the first week, mice were exposed to increasing concentrations of alcohol (*w/v*) starting with 3%, 6%, and 10% before being introduced to 20% alcohol for the rest of the paradigm. Fluid intake was measured in milliliters to calculate the g/kg intake and preference (%). Control groups that received only water bottles were run concurrently with the alcohol exposed groups. Bottle position on cages was switched each session and counterbalanced throughout the groups to ensure place preference was not established. Mice were split into high or low drinking groups based on a mean split of the total average drinking across all three weeks of the IA paradigm. Those above the mean were designated as high drinkers, and those below the mean were designated as low drinkers.

### DIDMSA paradigm

The Drinking-in-the-Dark-Multiple Scheduled Access (DID-MSA) procedure was performed as previously described (Melon, 2019) and adapted from (Bell et al., 2011; Melón et al., 2013). Briefly, beginning at lights out, mice are given access to water or a 20% unsweetened alcohol solution (95% ethanol Pharmco Products Inc., Brook-field, CT) during three, 1 hour drinking sessions, each separated by 2 hour breaks where only water was provided. Volumes consumed were recorded for each 1 hour binge-drinking session and summed for the day.

### Chronic Intermittent Ethanol (CIE) paradigm

The CIE paradigm was performed in vapor chambers acquired from La Jolla Alcohol Research Institute. Animals were randomly assigned to either vapor (CIE) or control (air) conditions. The target range of vapor chambers was 0.180-0.200 dL/mL. For two weeks Monday through Thursday, animals were placed in chambers for 16hrs (5:00pm to 9:00am). Animals were weighed the first day of each week. All animals received pyrazole (1mmol/kg) via intraperitoneal (IP) injections and CIE mice received a loading dose of ethanol IP (1.6g/kg). All animals received 0.3mL subcutaneous 0.9% saline. After removal from chambers, animals were tested on rotarod for intoxication levels and received 0.3mL subcutaneous saline. They recovered for eight hours and were abstinent Friday through Monday. CIE animals received hydrogel and nutrient gel and heat pad for post-ethanol recovery as well as during abstinence days. Submandibular blood was collected from one CIE and one air mouse on Thursday each week and plasma isolated for measurement of BEC.

### Spectral Analysis

Spectral analysis of the LFP/EEG signal was performed using the Short Time Fourier Transform (STFT) with a 5 second Hann window and 50% overlap. The power line noise (59 - 61 Hz) was removed and values were filled using nearest interpolation. Outliers in each spectrogram were identified using a two-stage process. Firstly, a time-series was obtained from the mean power across frequencies of each spectrogram. Large deviations, defined as those greater than the mean plus 4 times the standard deviation, were replaced with the median of the non-outlying data. Then, a sliding-window method was applied to detect more subtle outliers based on local context, using 5x the Median Absolute Deviation (MAD) of each 5-minute window. Resulting outliers were removed and replaced with forward fill interpolation of the nearest values. All resulting time-series, obtained from the mean power across frequencies, were manually inspected to remove bad regions that were replaced with median values for each frequency. Any resulting power spectral densities (PSDs) with no apparent peaks were rejected and were not included for further analysis. Each PSD was normalized to its total power across all frequencies analyzed (1 - 120 Hz). Then a value for each frequency band was obtained across all binge and break sessions. The normalized power for each frequency band and brain region were converted to z-scores.

### Phase-locking Analysis

To calculate the consistency of phase differences and measure coherence we calculated the PLV (Lachaux et al, 1999). Briefly, data were decimated to 250 Hz (FIR filter) and were then filtered across frequency bands using a Butterworth bandpass filter (order = 2). The instantaneous phases for BLA and FC channels were obtained by applying the Hilbert transform for each frequency band and the PLV was calculated by summing the phase differences (transformed into complex numbers) and then dividing their total magnitude by the length of the instantaneous phase (Lachaux et al, 1999). Higher PLV values indicate a stronger phase synchrony between the two regions. Only the signal from PSDs with clear peaks indicating a pronounced oscillation was included for PLV analysis.

### Elevated Plus Maze

All mice that received IA2BC were assessed for anxiety-like behavior in withdrawal six to eight hours after the last exposure to alcohol using the elevated plus maze (EPM). Mice were habituated in the EPM room for at least an hour before testing. The EPM apparatus consists of two 38 cm x 6.5 cm wide arms in a plus shape elevated 75 cm off the ground. Two arms are closed with 10 cm high walls and all arms have 48 equally spaced photocells. Mice were placed in the center of the maze to start the ten-minute test. The number of entries, distance traveled, and total time spent in the open and closed arms were measured using an automated software, Ethovision (Noldus).

### Immunofluorescence Staining

Immunofluorescence (IF) staining was performed as previously reported (Melón et al., 2018; DiLeo, 2022). For IA experiments, brain tissue was collected 24 hours after the last exposure to alcohol the day after anxiety-like behavior testing on the EPM. Mice were anesthetized with isoflurane, transcardially perfused with 0.9% saline, and 4% paraformaldehyde (PFA). The brains were rapidly excised, fixed in 4% PFA overnight, and subsequently cryoprotected in 10% and 30% sucrose. The brains were then flash-frozen using isopentane and stored at -80°C until cryosectioning on a Leica microtome. Free-floating 40 μm coronal slices were co-stained for PV and δ using universal antigen retrieval buffer (R&D systems CTS015) and primary antibodies against δ-GABA_A_R (1:100, Phosphosolutions 868A-GDN) and PV (1:1000, Sigma P3088) for 72 hours at 4°C. The slices were then incubated with a biotinylated goat anti-rabbit (1:1000, Vector Laboratories BA1000) and Alexa-Fluor 647 conjugated goat anti-mouse (1:200, ThermoFisher Scientific A28181) for two hours at room temperature and then streptavidin-conjugated Alexa-Fluor 488 (1:200, ThermoFisher Scientific S32354) for two hours at room temperature. Slices were mounted and coverslipped with antifade hard-set mounting medium with DAPI (Vectashield H1500).

### Histological Analysis

Fluorescent labeling in the BLA was imaged on a Nikon A1R confocal microscope and z-stacks were acquired using a 20X objective. Camera settings were kept consistent across samples and cohorts. The images were analyzed using Image J software by outlining PV-positive interneurons using the region of interest (ROI) manager and measuring the integrated density of PV and δ expression on the outlined PV-positive interneurons. Each cell was considered its own data point within each animal.

### Statistical Analysis

Data were analyzed using Prism software (GraphPad), MATLAB (MathWorks) and Python. To ensure a consistent time period for analysis across cohorts, we analyzed the first 40 minutes of baseline and the first 35 minutes of each injection period. For analysis of effects on LFP frequencies, repeated measures of two-way ANOVAs were performed to detect the significance of frequency, treatment, sex, or genotype. For analysis of IF data after IA, a one-way ANOVA was used to detect significant differences between drinking groups. A Greenhouse-Geisser correction was applied where necessary. A mixed-effects model was used if values were missing across days. A post hoc Šídák’s multiple comparisons test was performed to identify significant differences in specific groups. *p* values < than 0.05 were considered significant. All *n* values for each treatment group are shown in the figure legends. For the DID-MSA data, boxplots were created in PRISM and significance was tested using two-way ANOVAs (frequency vs session) followed by multiple-t-test adjusted using the Sidak method. Scatter plots with regression line were created using the seaborn lmplot. Correlation was assessed using a Pearson correlation and significance was assessed using a permutation of the shuffled values (N = 10000 permutations). Resulting p-values were corrected for multiple comparisons using false discovery rate (Benjamini/Hochberg method).

## Results

### Intermittent Access Escalates Alcohol Intake in C57BL/6J mice

Variation in alcohol drinking exists across strains, species, and sex from rodents to humans (Agabio, Pisanu, Gessa, & Franconi, 2016; Hwa et al., 2011; Ritchie & Roser, 2018; Sneddon, White, & Radke, 2019; Yoneyama, Crabbe, Ford, Murillo, & Finn, 2008). Although there is evidence of genetic vulnerability to high levels of alcohol drinking in humans, this does not address the additional environmental influences that contribute to an individual’s risk for developing an AUD (Crabbe, Phillips, & Belknap, 2010; Enoch, 2003). Therefore, efforts to understand vulnerability or susceptibility to high alcohol intake are important in order to identify or predict individuals who may go on to develop an AUD. Since we determined that acute alcohol could modulate BLA network states in C57BL/6J mice (DiLeo, 2022) and we have shown that disturbed network states can influence mood disorders and behavior (Antonoudiou et al., 2022; Walton et al., 2022), we hypothesized that the BLA network response to acute alcohol could predict or correlate with voluntary alcohol intake and withdrawal-induced anxiety-like behaviors. To test this hypothesis, we recorded local field potentials (LFP) in the BLA of male and female C57BL/6J mice during acute alcohol exposure (Figure 1) and a three-week, IA2BC procedure (Figure 1A). IA2BC escalated alcohol intake, but not water intake, from session 1 to session 9 in males (*p* = 0.0001, 95% C.I = [4.013, 13.06]) and females (*p* < 0.0001, 95% C.I = [16.77, 46.17]; (Figure 1B-C). Females (*M* = 14.86, *SEM* = 1.319) had a higher average alcohol intake over three weeks compared to males (*M* = 7.37, *SEM* = 0.854; *t* (40) = 4.419, *p* < 0.0001; (Figure 1D), a finding which is well established for this and other voluntary drinking paradigms (Hwa et al., 2011). Preference for alcohol did not significantly change from sessions 1 to 9, and there were no differences between males and females (Figure 1E). Total fluid intake and body weights were not different between water and alcohol groups for male (Figure 1-1A-B) or female C57BL/6J mice (Figure 1-1C-D).

**Figure 1:**
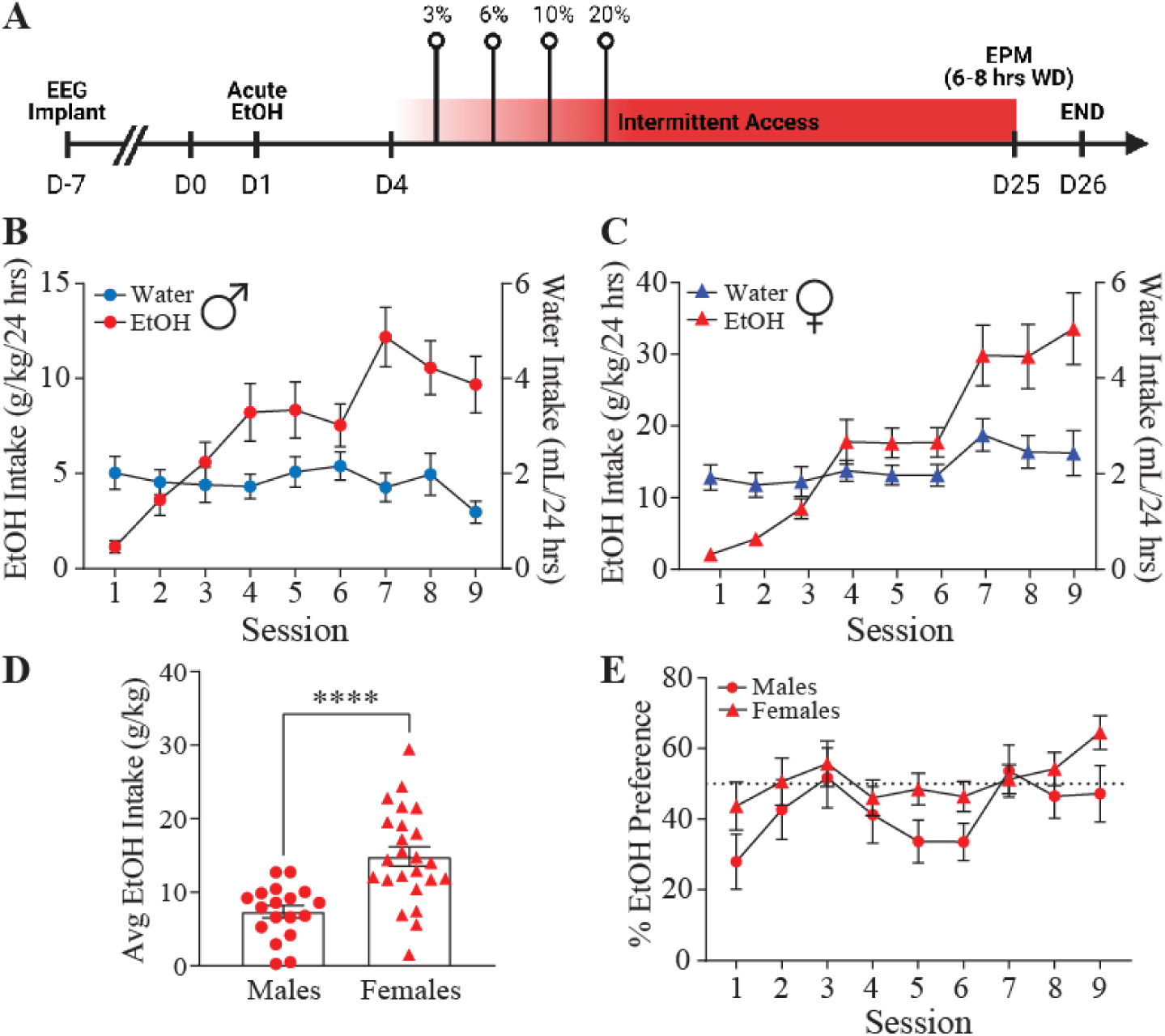
Intermittent Access Escalates Alcohol Intake in C57BL/6J mice. A, A schematic of the experimental design with a timeline of LFP recording in response to acute ethanol exposure preceding the three-week, IA2BC procedure and subsequent behavioral assessments. The average water (blue) and ethanol (red) consumed per session throughout the IA2BC paradigm in males (B; *n* = 18) and females (C; *n* = 24). D, The average alcohol intake consumed during the IA2BC paradigm is higher in females than in males with no difference in alcohol preference (E). Individual and mean ± SEM values are graphed. Statistical analysis by unpaired t-test (D). \*\*\*\**p* < 0.0001 vs females.

To assess anxiety-like behavior associated with alcohol intake, we tested all mice in the EPM six to eight hours after the last exposure to alcohol. We found that alcohol exposed female C57BL/6J mice spent more time in the open arms (p = 0.001, 95% C.I = [41.71, 139]) and less time in the closed arms of the EPM (p = 0.003, 95% C.I = [-134.4, -37.15]; Figure 2D) and traveled an increased distance in the open arm of the EPM (p = 0.013, 95% C.I = [63.73, 618]; Figure 2F) without a change in the number of entries (Figure 2E). This data suggests that alcohol exposed female mice exhibited less anxiety-like behavior in this test compared to water controls, which may be due to the temporal relationship between alcohol exposure and behavioral testing. There were no differences in the performance in the EPM in males exposed to alcohol or water (Figure 2A-C). Lastly, there was no difference between low and high drinking females in any of the EPM measures (data not shown).

**Figure 2:**
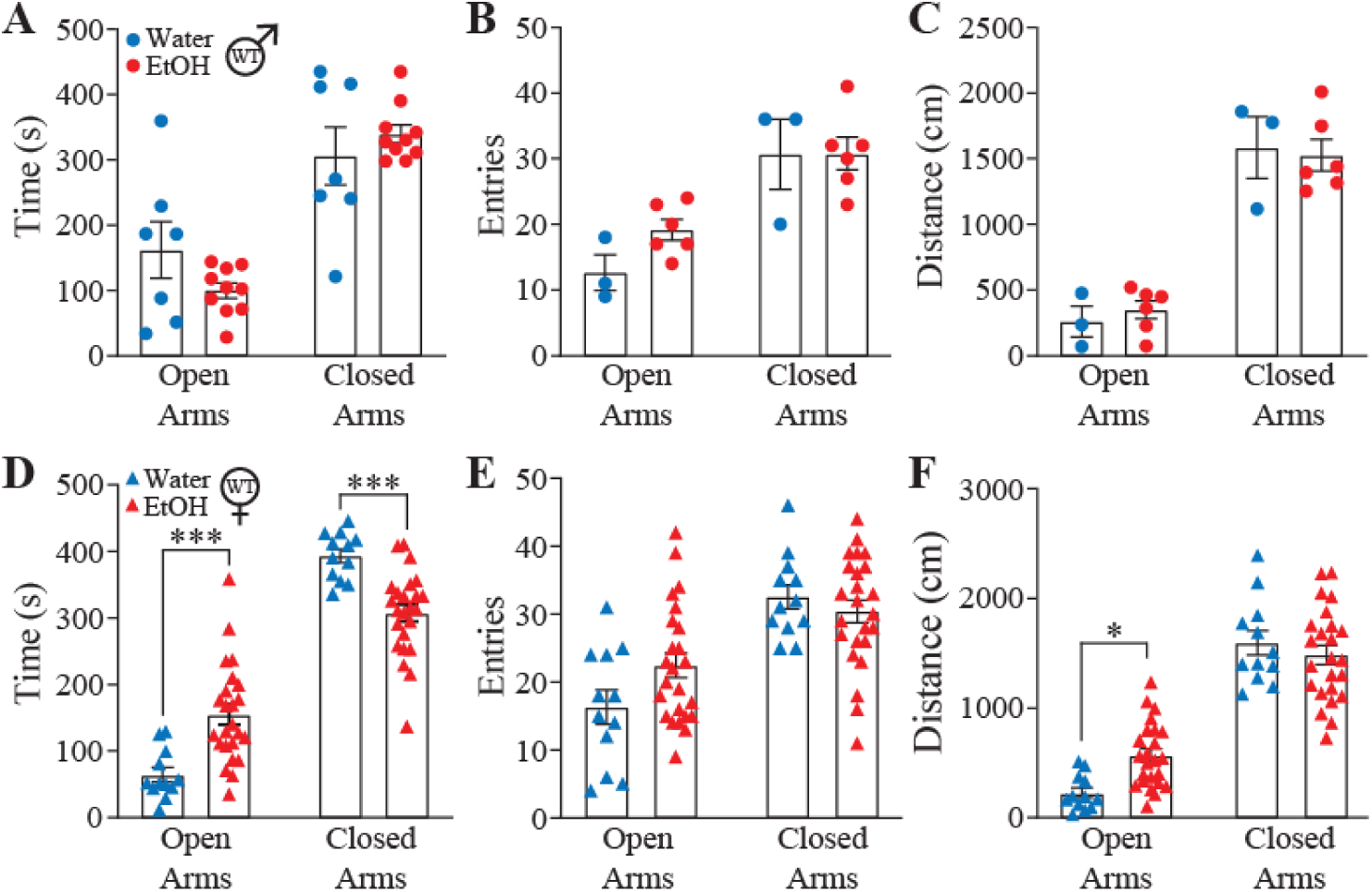
Sex differences in avoidance behaviors following alcohol exposure. Avoidance behaviors were assessed 6-8 hrs following the IA2BC paradigm. The average amount of time (A, D), number of entries (B, E), and total distance traveled (C, F) in the open arm of the elevated plus maze in males (A-C; water *n* = 7; EtOH *n* = 9) and females (D-F; water *n* = 12; EtOH *n* = 24). Females exposed to alcohol exhibit a decrease in avoidance behaviors in the elevated plus maze compared to water drinkers (D). Individual and mean ± SEM values are graphed. Statistical analysis by two-way ANOVA. \**p* < 0.05, \*\*\**p* < 0.005 vs water. Note: *n* number for males (water *n* = 3; EtOH *n* = 6) differ for number of entries and distance moved in EPM.

### BLA Response to Acute Alcohol Correlates with Voluntary Drinking Levels

To understand if there was a relationship between the BLA LFP response to acute alcohol and voluntary drinking levels, we split the alcohol groups into high or low drinkers by the mean alcohol intake across the three weeks of drinking. Those above the mean were designated as high drinkers and those below the mean were designated as low drinkers. When the groups were split, there were significant differences between the average drinking levels and preference in males (*M* = 7.309, *SEM* = 1.115; Figure 3A,B) and females (*M* = 17.89, *SEM* = 3.822; *t* (16) = 2.658, *p* = 0.017; Figure 3C,D).

**Figure 3:**
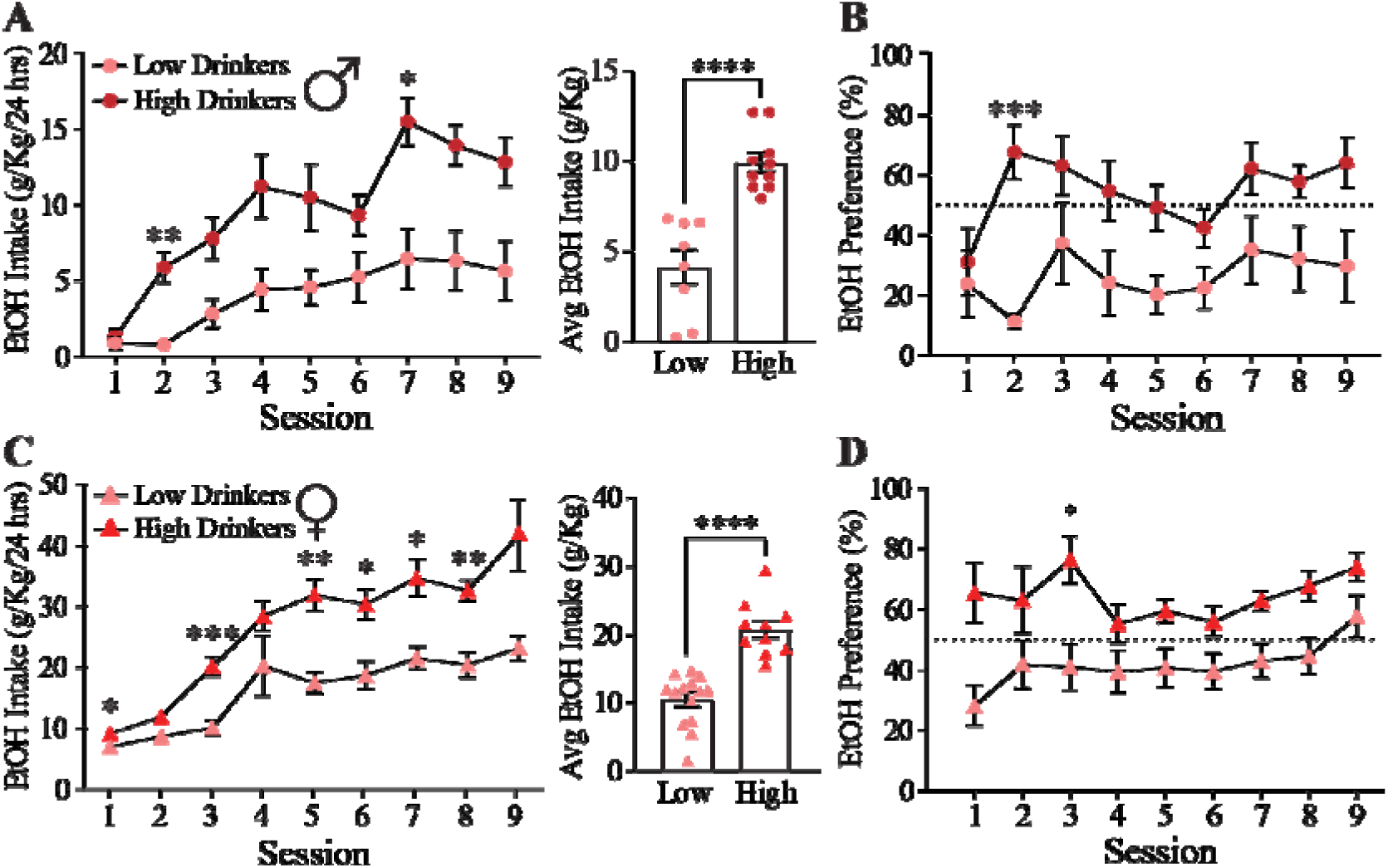
Identification of high and low alcohol drinkers in the IA2BC paradigm. The mean alcohol intake across the three weeks of drinking was used to split animals into high (above the mean) and low (below the mean) drinking groups. Average ethanol intake per session and total is significantly increased in the high drinking group in males (A; low *n* = 8; high *n* = 10) and females (C; low *n* = 14; high *n* = 10) compared to the low drinking group with an increase in ethanol preference (B, D). Individual and mean ± SEM values are graphed. Statistical analysis by two-way ANOVA. \**p* < 0.05, \*\**p* < 0.01, \*\*\**p* < 0.005, \*\*\*\**p* < 0.0001.

We then analyzed the relationship between BLA oscillatory states in response to acute alcohol exposure and found that alcohol reduced low and high gamma in both male and female C57BL/6J mice (Figure 4A-B), similar to previous findings (DiLeo, 2022). We did not find any differences between the frequency of oscillations in the BLA at baseline, during the vehicle injection, during the alcohol injection, or in the change between periods in low or high drinking groups (Figure 4-1). We also did not find any significant differences between low and high drinking groups when we grouped all mice together (Figure 4C). However, there is a significant negative correlation between the average alcohol intake and the normalized γ power during exposure to acute alcohol (*r* (35) = -0.35, *p* = 0.035; Figure 4D), demonstrating that the animals in which acute alcohol exposure least impacted BLA network activity consumed more alcohol. Further, we demonstrate that both high and low drinkers exhibit an increased amount of time and distance traveled in the open arm of the EPM compared to water drinkers (Figure 4-2). These data demonstrate a relationship between the ability of acute alcohol to modulate BLA network states, the extent of voluntary alcohol consumption, and the anxiolytic effects of alcohol.

**Figure 4:**
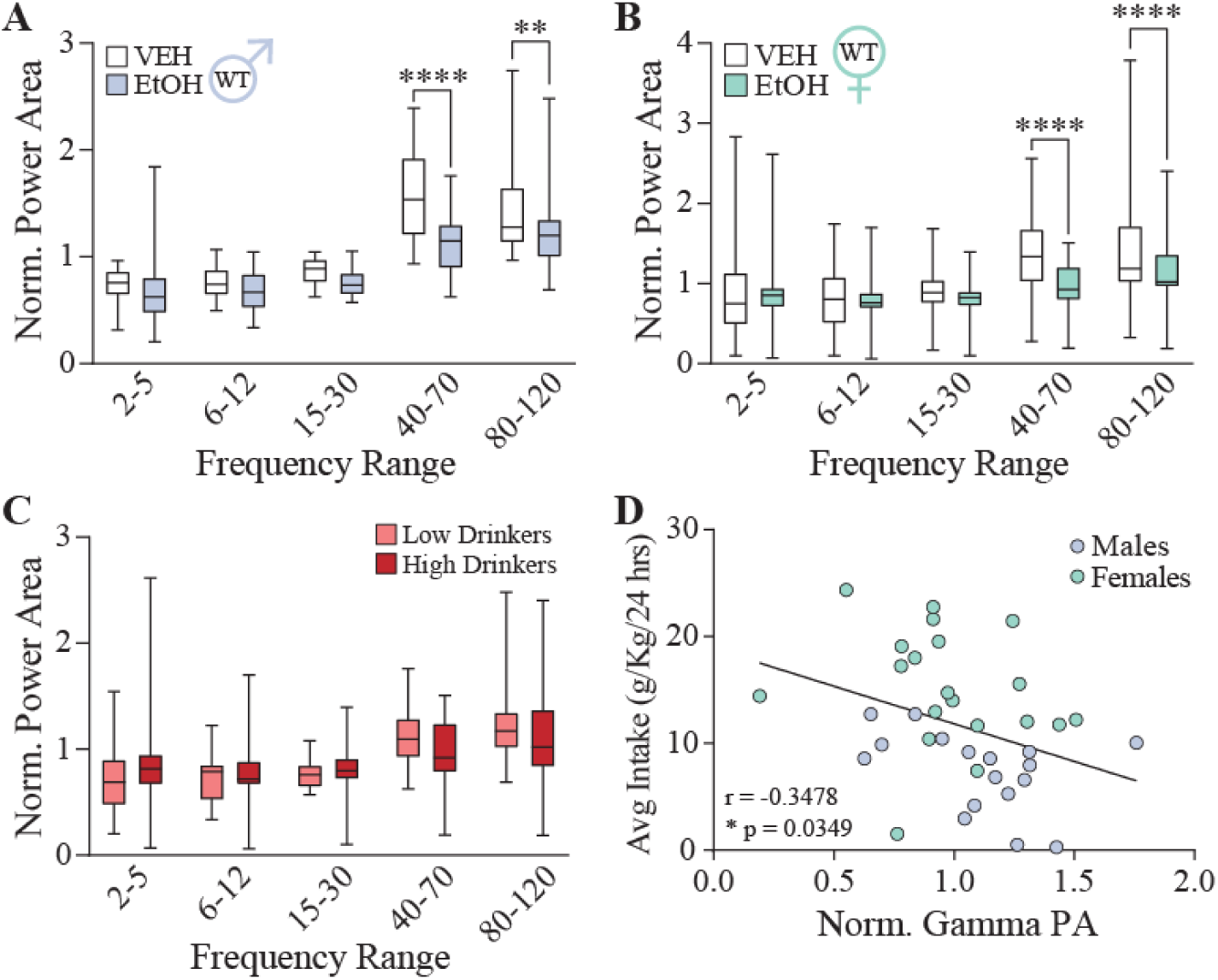
Gamma oscillations in the BLA predict the extent of voluntary alcohol consumption. Acute ethanol exposure decreases the power of gamma oscillations in the BLA in both males (A; *n* = 17) and females (B; *n* = 20). The average power across frequencies is not different between high and low drinking groups (C). D, There is a significant negative correlation between the ability of acute alcohol to modulate BLA gamma oscillations and the subsequent amount of voluntary alcohol consumed (*r* (35) = -0.35, *p* = 0.035). Statistical analysis by two-way ANOVA (A-C) and correlation (D). \*\**p* < 0.01, \*\*\*\**p* < 0.0001.

### Intermittent Access Differentially Modulates δ-GABA_A_R expression in the BLA

Lastly, due to the well-established sex differences in alcohol intake in voluntary drinking paradigms and in the expression of the δ-GABA_A_Rs (DiLeo, 2022), we measured the expression of δ -GABA_A_Rs on PV interneurons in the BLA. We found that low and high drinking animals had increased PV immunoreactivity (low: *p* < 0.0001, 95% C.I = [-149223 to -69630]; high: *p* < 0.0001, 95% C.I = [-129562 to -44624]) and expression of δ-GABA_A_Rs on PV interneurons (low: *p* = 0.001, 95% C.I = [-75305, -18905]; high: *p* = 0.001, 95% C.I = [-78793, -18604]) in male C57BL/6J mice compared to water drinking mice (Figure 5A-B left panels).

**Figure 5:**
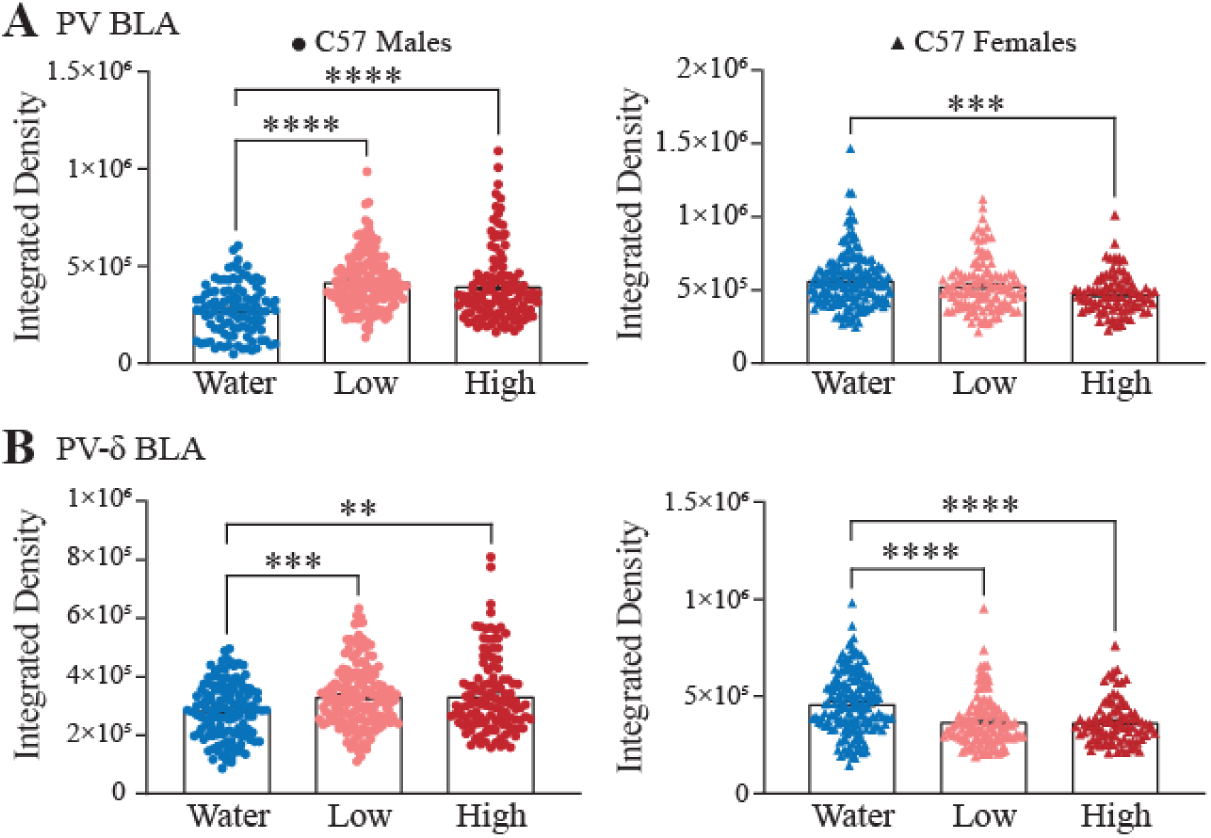
Voluntary ethanol consumption differentially modulates δ-GABA_A_R expression in the BLA. A, The average parvalbumin expression in the BLA is increased in males (water: cell *n* = 137, animal *n* = 4; low: cell *n* = 166, animal *n* = 3; high: cell *n* = 127, animal *n* = 3) and females (water: cell *n* = 171, animal *n* = 4; low: cell *n* = 119, animal *n* = 4; high: cell *n* = 90, animal *n* = 4) following the IA2BC paradigm. B, GABA_A_R δ subunit expression on PV interneurons in the BLA is increased in both high and low drinking male mice, whereas GABA_A_R δ subunit expression on PV interneurons in the BLA is decreased in both high and low drinking female mice. Statistical analysis by one-way ANOVA. \*\**p* < 0.01, \*\*\**p* < 0.005, \*\*\*\**p* < 0.0001 vs water.

Interestingly, we found the opposite effect in female mice. PV immunoreactivity was reduced in high drinking females (*p* < 0.0001, 95% C.I = [43789, 143418]) with both low (*p* < 0.0001, 95% C.I = [56951, 130212]) and high (*p* < 0.0001, 95% C.I = [55960, 135878]) drinking groups having reduced δ-GABA_A_R compared to water drinkers (Figure 5A-B, right panels). These data demonstrate a relationship between δ-GABAAR expression and voluntary alcohol intake.

### Altered BLA Network States Translates to Other Models of Alcohol Exposure and Withdrawal

To better appreciate the impact of voluntary alcohol intake on BLA network states, we employed the Drinking in the Dark Multiple Scheduled Access (DID-MSA) paradigm (Figure 6A). The power of oscillations across frequency bands is quantified across binge and break periods throughout the DID-MSA protocol in the BLA and frontal cortex and the coherence across regions is measured (Figure 6-1). While there were no significant differences in the power of the oscillations across frequencies in either the BLA (F(2.024, 12.14) = 0.8129, p = 0.47, n = 7 (3 male, 4 female), Figure 6B) or frontal cortex (F(1.281, 6.403) = 0.1608, p = 0.76, n = 6 (2 male, 4 female), Figure 6C) and no change in the coherence between the BLA and frontal cortex (F(2.162, 10.81) = 1.361, p = 0.30, n = 6 (2 male, 4 female), Figure 6D), we did observe a relationship between oscillation power and the extent of ethanol intake during binge sessions (Figures E-F). The power of oscillations within the low theta frequency (2-5 Hz) in the BLA (r = -0.26, p = 0.02, n = 95 sessions (11 mice), Figure E, top) and frontal cortex (r = -0.37, p < 0.0001, n = 89 sessions (11 mice), Figure F, top) was inversely correlated with the extent of alcohol intake. In contrast, the power of gamma oscillations (40-70 Hz) was positively correlated with the amount of ethanol intake in both the BLA (r = -0.28, p = 0.003, n = 95 sessions (11 mice), Figure E, bottom) and frontal cortex (r = -0.42, p < 0.0001, n = 89 sessions (11 mice), Figure F, bottom). These data demonstrate that the more alcohol voluntary consumed the greater the modulation of BLA network activity. There was no effect of ethanol consumption on coherence between BLA and frontal cortex (p > 0.05, Figure 6-2, bottom-row). These findings demonstrate that alcohol influences BLA and frontal cortex network states in the DID-MSA paradigm in relation to the extent of voluntary alcohol consumption.

**Figure 6:**
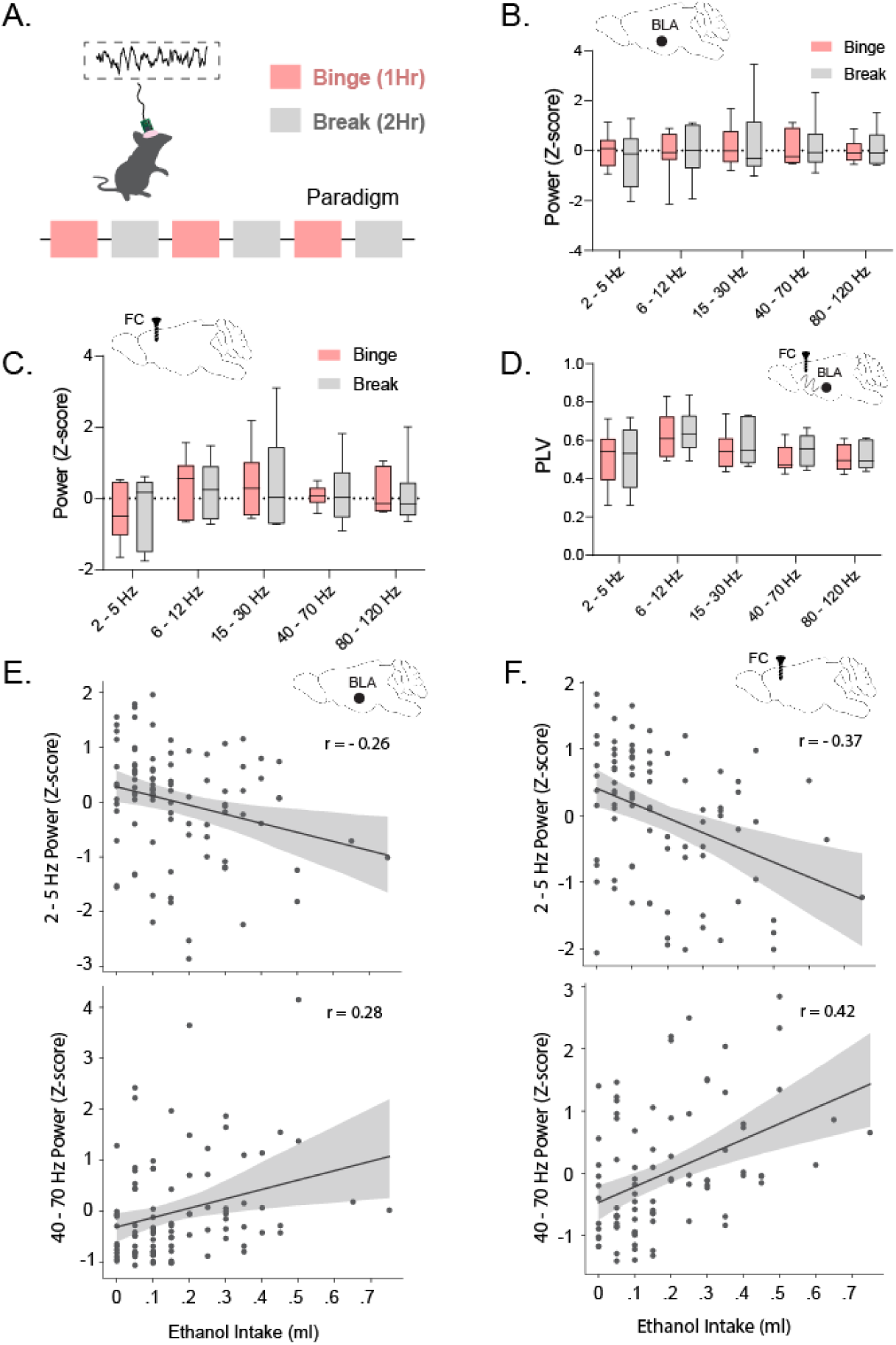
Binge drinking alters oscillatory states in the BLA. A, A schematic of the DID-MSA paradigm. The average power of oscillations across frequencies in the BLA (n = 7 (3 male, 4 female)), (B), frontal cortex (n = 6 (2 male, 4 female)), (C), and coherence between the BLA and frontal cortex (n = 6 (2 male, 4 female)) (D) during binge and break periods of the DID-MSA paradigm. There is a negative correlation between oscillatory states in the BLA (n = 95 sessions (11 mice)), (E) and frontal cortex (n = 89 sessions (11 mice)) (F) in the theta range (E, F, top) and a positive correlation between oscillations in the gamma range (E, F, bottom) and the amount of ethanol consumed in the DID-MSA binge periods. Statistical analysis in box plots by two-way repeated measures ANOVA and in correlations plots by Pearson correlation with permutation (see *Statistical Analysis* section).

To examine the impact on alcohol dependence and withdrawal on BLA network states, we utilized the CIE paradigm (Figure 7A). Unlike the impact of alcohol exposure on BLA network states, which is associated with an increase in γ power, alcohol withdrawal following CIE exposure is associated with a decrease in γ power in the BLA (Figure 7B-C; 2-way ANOVA: F 4, 48) = 2.568, p=0.045; for γ power: Sidak’s test with p = 0.029), but not the fontal cortex (Figure 7B,D; 2-way ANOVA: F(4, 52) = 1.848, p=0.134), compared to air exposed controls. Collectively, these data demonstrate that voluntary alcohol exposure modulates BLA network activity and withdrawal is associated with opposing effects on BLA network states.

**Figure 7:**
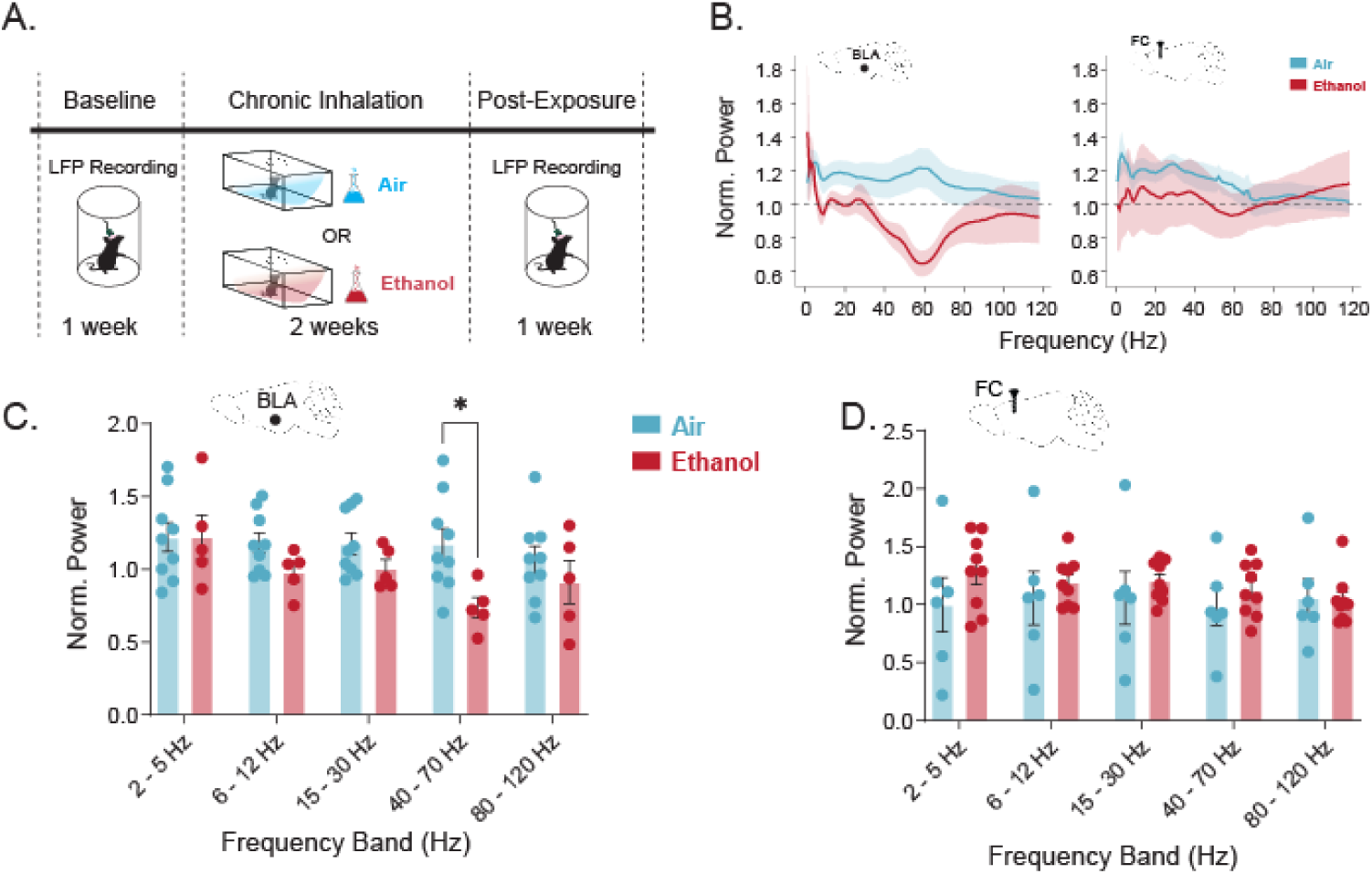
CIE reduces γ-band oscillations in the BLA. (A) Schematic of the CIE paradigm. LFP recordings were performed before and after chronic inhalation of air or ethanol. (B) Power spectral density plots normalized to baseline period in BLA (air: n = 9, ethanol: n = 5) and FC (air: n = 9, ethanol: n = 6), where shaded regions indicate SEM. Summary bar plots of normalized power in (C) BLA and (D) FC, where dots represent individual animals and error bars represent SEM. Statistical analysis by two-way ANOVA followed by Sidak’s multiple comparisons tests. \**p* < 0.05.

## Discussion

In order to test the hypothesis that oscillatory states in the BLA in response to acute alcohol could predict or correlate with voluntary alcohol drinking, we recorded network activity in the BLA during an acute alcohol paradigm combined with an IA2BC paradigm for three weeks to escalate alcohol intake in male and female C57BL/6J mice. We also examined the relationship between BLA oscillatory states and alcohol consumption in the DID-MSA paradigm. We observed that acute alcohol exposure decreased γ power in both males and females, which is consistent with results found by others (Henricks et al., 2019). This may indicate that alcohol is disrupting the rhythmicity or synchronicity of pyramidal cells by influencing PV interneuron inhibition (Antonoudiou et al 2022). Importantly, these effects of alcohol on oscillations in the BLA are unique from other GABA modulators, including diazepam and allopregnanolone (Antonoudiou et al 2022). Although we did not find significant differences in the average power of BLA network states at baseline between low and high drinking animals, we did find a significant correlation between average drinking levels and γ frequency in response to acute alcohol. It is possible that the number of animals we had designated in low and high drinking groups was too low to detect any differences and the correlation between LFP frequency and drinking was only detected when male and female mice were analyzed together. Additionally, the IA2BC paradigm is traditionally used for six to eight weeks, and it is possible that the levels of alcohol intake were still escalating at three weeks. Future studies could extend the length of the IA2BC paradigm or use a paradigm with a shorter period of alcohol access, like DID, to facilitate higher alcohol intake that may make it easier to detect changes to network states. In fact, we utilized the DID-MSA paradigm and demonstrated that voluntary alcohol consumption in this paradigm was capable of altering BLA network states which were proportional to the extent of alcohol consumption. Further, we demonstrate that alcohol withdrawal is associated with opposing changes on the BLA network state to those observed with alcohol exposure, further suggesting that BLA network states are influenced by alcohol.

Interestingly, we did not find sex differences in alcohol’s effects on BLA network activity even though male and female mice have differing sensitivity to the rewarding and aversive properties of alcohol (Flores-Bonilla & Richardson, 2020; Hwa et al., 2011; Vetter-O’Hagen, Varlinskaya, & Spear, 2009). Sex differences have been reported in neural oscillations in major depressive disorder and rodent models of post-traumatic stress disorder, suggesting we may see similar differences in responses to alcohol (Anker & Carroll, 2011; Fenton et al., 2016). Sex differences in alcohol related anxiety-like behavior is also well-documented (Barkley-Levenson & Crabbe, 2015; Rhodes et al., 2005) and may be related to altered BLA network states given the relationship to behavioral states (Antonoudiou et al., 2022; Davis et al., 2017; Likhtik et al., 2013; Ozawa et al., 2020; Stujenske, Likhtik, Topiwala, & Gordon, 2014). Additionally, sex differences have been reported in neural oscillations in major depressive disorder with oscillatory signatures of susceptibility (Theriault, Manduca, & Perreault, 2019).

After three weeks of IA2BC, we assessed all mice for anxiety-like behavior using the EPM six to eight hours after the last exposure to alcohol. We did not find any differences between male alcohol exposed and water control mice on the EPM. However, we did find that alcohol exposed female C57BL/6J mice spent more time and traveled more on the open arm compared to water controls, indicative of less anxiety-like behavior. This could be interpreted as a degree of risk-taking behavior where alcohol exposed mice are willing to take a risk in a novel environment to seek drug. The increased distance traveled could also suggest hyperlocomotion in these animals, which makes it difficult to interpret these results. Others have reported both hypo and hyper locomotion of alcohol exposed females on the EPM (Albrechet-Souza, Schratz, & Gilpin, 2020; Holleran & Winder, 2017; Kliethermes, 2005), making it difficult to interpret anxiety-like behavior in this assay. It is also possible the three weeks of IA2BC was not long enough to induce anxiety-like behaviors in withdrawal. Others have reported depressive-like behaviors in the forced swim test and novelty suppressed feeding test, but not anxiety-like behavior in the EPM or elevated zero maze using longer paradigms (Bloch, Rinker, Marcus, & Mulholland, 2020; Holleran & Winder, 2017; Holleran et al., 2016; Pang, Renoir, Du, Lawrence, & Hannan, 2013). This suggests that additional tests may be required to assess affective behavior in withdrawal or the ideal length of time into withdrawal to capture these effects is difficult to resolve. Further, alcohol naïve female C57BL/6J mice show less anxiety-like behavior in these assays compared to males, indicating that the mice are starting off with different baseline behaviors that may interact with alcohol in different ways (Albrechet-Souza, Schratz, & Gilpin 2020; Bloch et al., 2020).

There are well-documented sex differences in responses to alcohol and estrous-cycle dependent δ expression (Barkley-Levenson & Crabbe, 2015; Maguire, Stell, Rafizadeh, & Mody, 2005; Rhodes, Best, Belknap, Finn, & Crabbe, 2005). Here we demonstrate sex differences in the expression of δ-GABA_A_Rs in the BLA, which may contribute to the differences in the level of voluntary alcohol consumption. Since δ-GABA_A_Rs are implicated in alcohol drinking behaviors (Darnieder et al., 2019; Melón, Nasman, John, Mbonu, & Maguire, 2018), we investigated whether levels of alcohol intake in IA2BC modulated δ expression. Indeed, we found that alcohol exposure, regardless of drinking level, increased PV immunoreactivity and δ expression on PV interneurons in male C57BL/6J mice while reducing δ expression in females. These data suggest that there is a relationship between the naïve BLA’s response to acute alcohol and future voluntary drinking, demonstrating a relationship between the ability of alcohol to modulate BLA network states and alcohol consumption. Although δ subunits are important players in carrying out alcohol’s effects, other GABAergic subunits expressed on PV cells may also be important to shifting oscillatory power. The γ2 subunit has also been implicated in anxiety-like behavior (Chandra, Korpi, Miralles, De Blas, & Homanics, 2005) and a genetic variant of γ2 is associated with alcohol withdrawal severity (Buck & Hood, 1998).

Given the heterogeneity of the interneuron population in the BLA, it is possible other interneuron types, like somatostatin, cholecystokinin, or PKC-δ expressing cells are involved (Klausberger et al., 2005). Influence of intercalated paracapsular cells surrounding the amygdala, GABAergic cells in the central amygdala, and other interneuron types in the BLA may also be involved in mediating effects of alcohol (Babaev, Piletti Chatain, & Krueger-Burg, 2018; Marowsky & Vogt, 2014; Roberto, Madamba, Stouffer, Parsons, & Siggins, 2004). However, the literature supports a strong role for PV interneurons in oscillation generation, giving support to the likely fact that alcohol’s effects on PV interneurons are influencing the oscillations (Antonoudiou et al., 2022). Additionally, because LFPs may be reflections of inputs to an area and evidence has shown that oscillations can originate in other places (Carmichael et al., 2017), it is possible that the effects seen in the BLA are from another region, like the mPFC (Janak & Tye, 2015; Koob & Volkow, 2009). However, we recently demonstrated that the BLA is capable of generating intrinsic oscillations, which involve PV interneurons (Antonoudiou et al., 2022).

Despite the extensive literature on the GABAmimetic effects of alcohol, the direct effects of alcohol on GABAergic neurotransmission are still unclear and controversial (Weiner & Valenzuela, 2006). It is possible that alcohol’s indirect effects on receptor expression, neurotransmitter availability, and other neuromodulators could account for the changes in BLA oscillations observed here (Fleming, Manis, & Morrow, 2009; Morrow, VanDoren, Penland, & Matthews, 2001; Olsen & Liang, 2017). Additional research is required to fully understand the mechanisms mediating the impact of alcohol on BLA network states. Despite these remaining questions, the current study nicely demonstrates the ability of alcohol to modulate BLA network states and predict future voluntary alcohol consumption which necessitates digging deeper into the mechanisms through which alcohol impacts BLA network states and the inherent differences in the responsivity of this network to the effects of alcohol. Further, we demonstrate opposing effects of alcohol withdrawal on BLA network states which we propose may contribute to negative affective states and hyperkatefia and drive to increase alcohol intake. Future studies are required to evaluate the relevance of these BLA network states to the cycle of AUDs.

## Acknowledgements

The authors would like to thank Dr. Klaus Miczek and Dr. Leon Reijmers for thoughtful feedback and guidance on this project.

---

**Figure 1-1:**
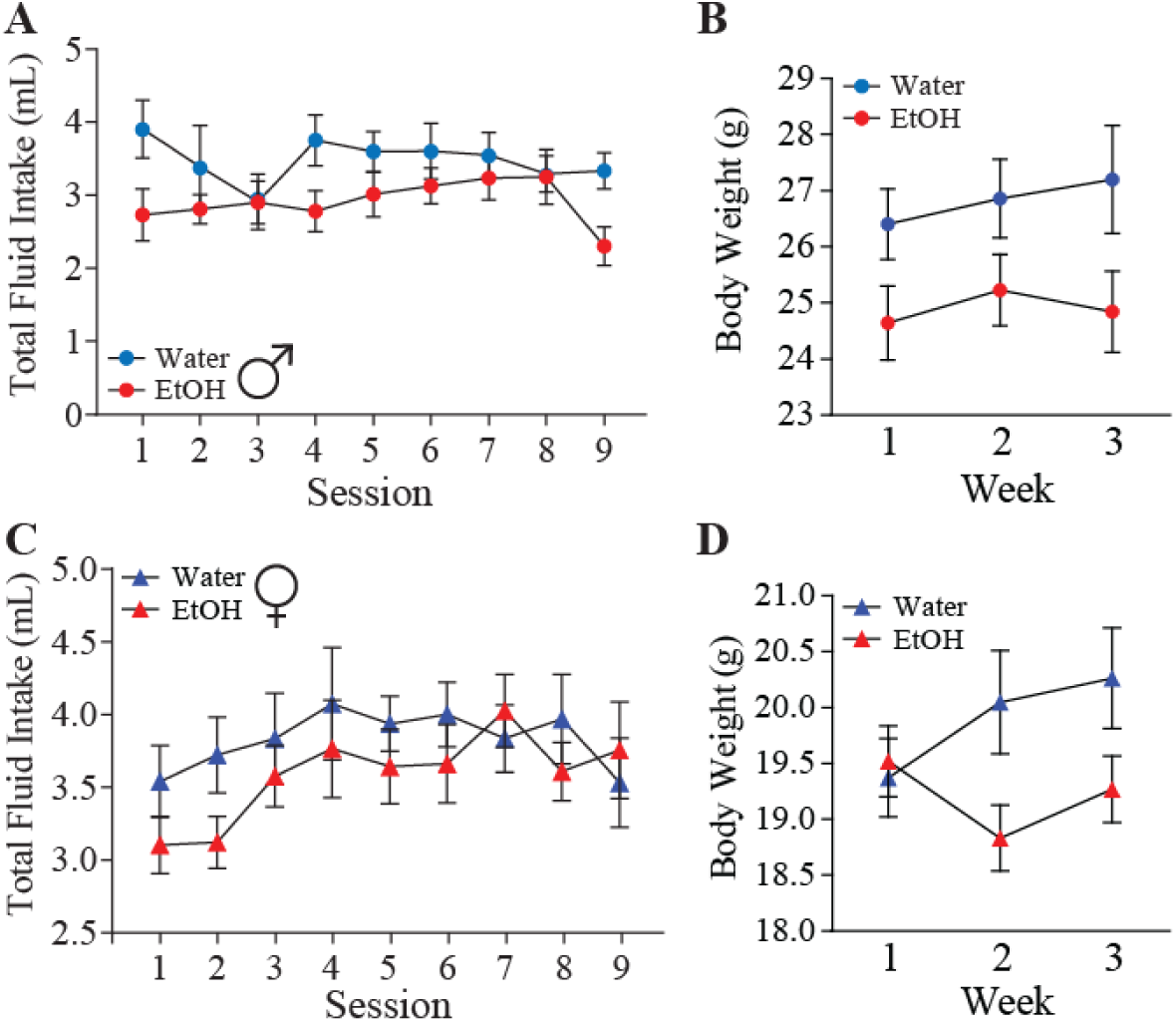
The IA2BC paradigm does not alter fluid intake or body weight. Total fluid intake (A, C) and body weights (B, D) in male (A-B; water *n* = 7; EtOH *n* = 18) and female (C-D; water *n* = 12; EtOH *n* = 24) water drinkers (blue) or ethanol exposed (red). Statistical analysis by two-way ANOVA.

**Figure 4-1:**
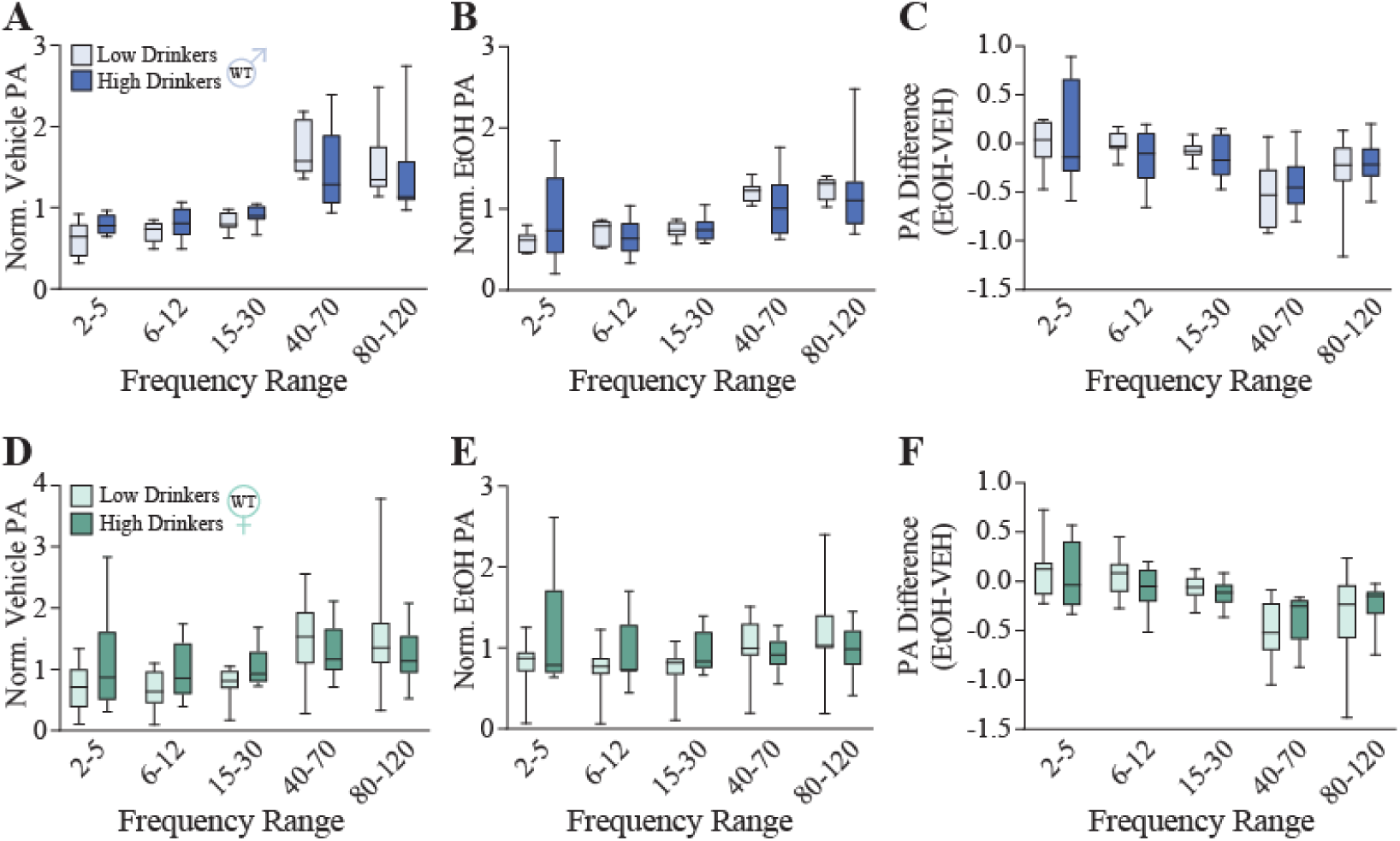
Alcohol modulation of BLA oscillatory states in high and low drinkers. The power of oscillations across frequencies in response to vehicle injection (A, D), acute ethanol exposure (B, E), or the difference between vehicle and ethanol injections (C, F) in high and low drinking males (A-C; low *n* = 7; high *n* = 10) and females (D-F; low *n* = 11; high *n* = 9). Statistical analysis by two-way ANOVA.

**Figure 4-2:**
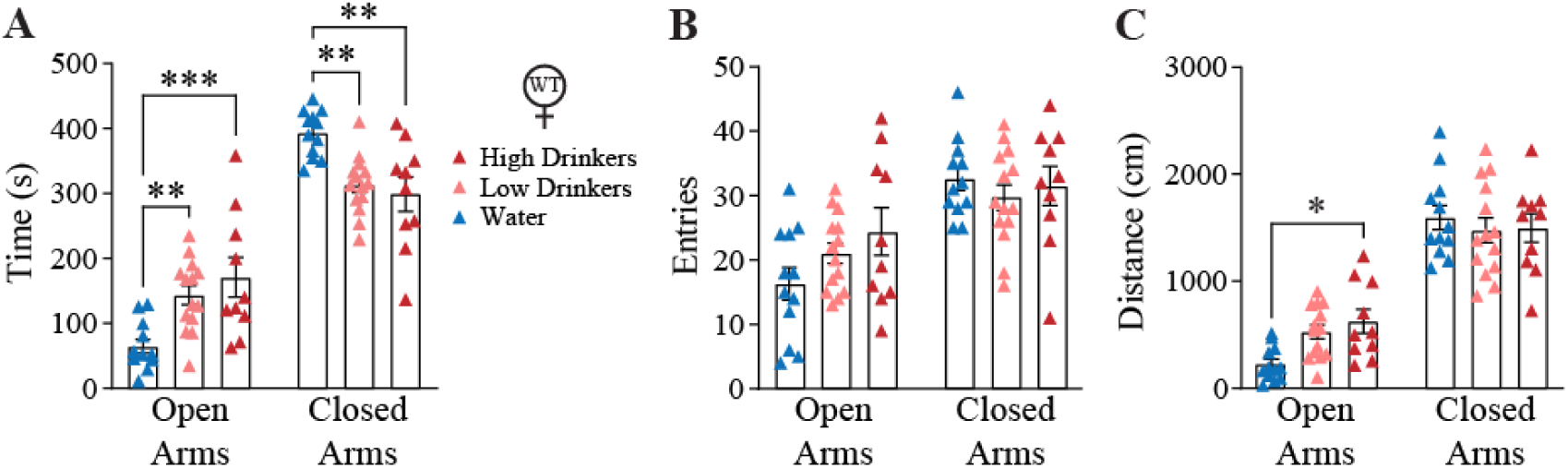
The influence of the amount of alcohol consumed on avoidance behaviors. Avoidance behaviors were assessed in high and low drinking females following the IA2BC paradigm. The average amount of time (A), number of entries (B), and total distance traveled (C) in the open arm of the elevated plus maze was assessed in high and low drinking females (water *n* = 12; low *n* = 14; high *n* = 10) . Females exposed to alcohol exhibit a decrease in avoidance behaviors in the elevated plus maze compared to water drinkers with no significant difference between high or low drinkers. Statistical analysis by two-way ANOVA. \**p* < 0.05, \*\**p* < 0.01, \*\*\**p* < 0.005, vs water.

**Figure 6-1:**
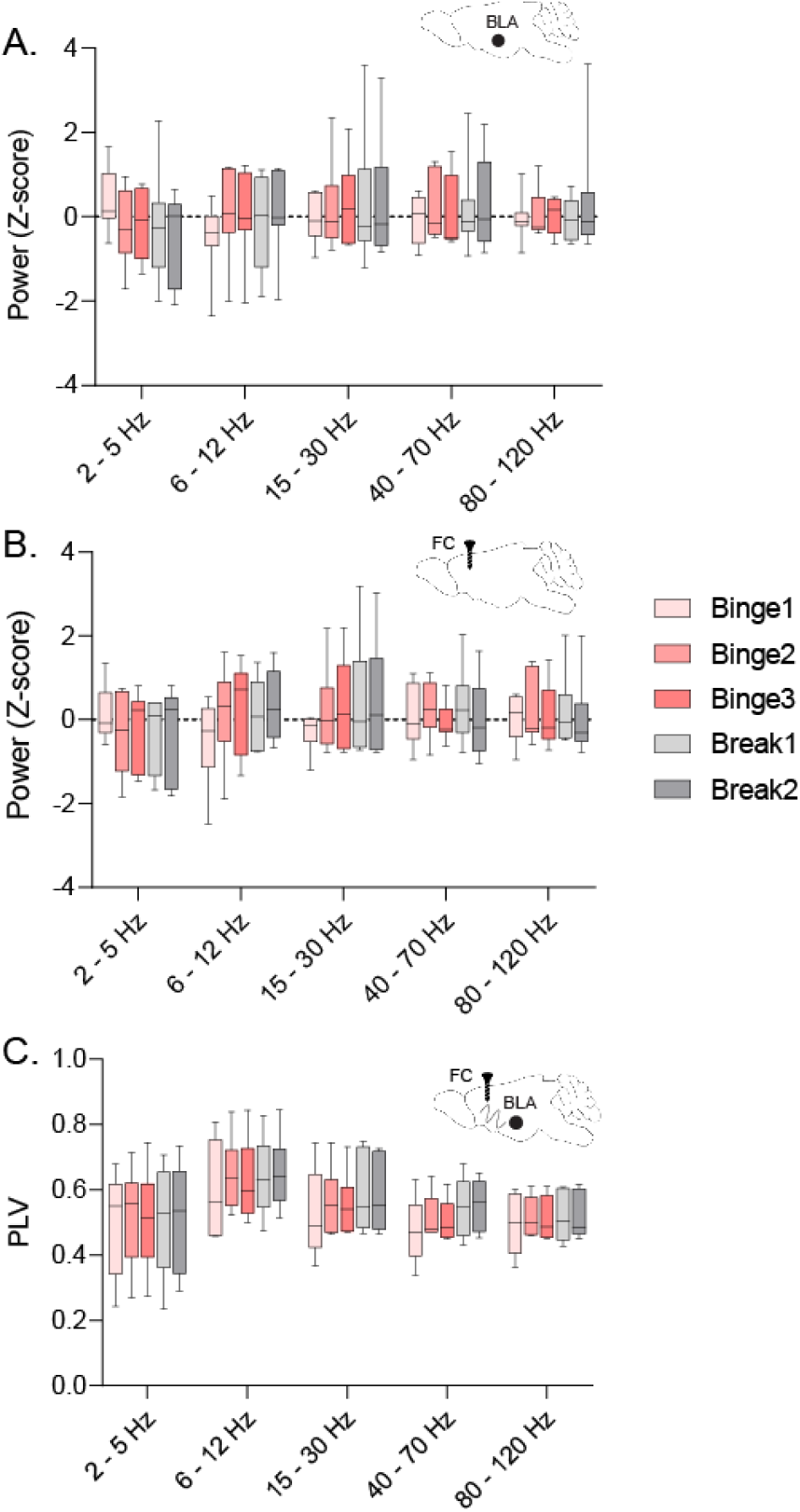
Oscillatory states during binge drinking. Oscillatory states during binge drinking. The average power of oscillations across frequencies in the BLA (F(3.221, 19.33) = 1.127, p=0.3656, n = 7 (3 male, 4 female)) (A), frontal cortex (F(16, 108) = 0.3853, p=0.9835, n = 7 (3 male, 4 female)) (B), and coherence between the BLA and frontal cortex (F(16, 96) = 0.2082, p=0.9995, n = 6 (2 male, 4 female)) (C) during individual binge and break sessions throughout the DID-MSA protocol. Statistical analysis by two-way repeated measures ANOVA.

**Figure 6-2:**
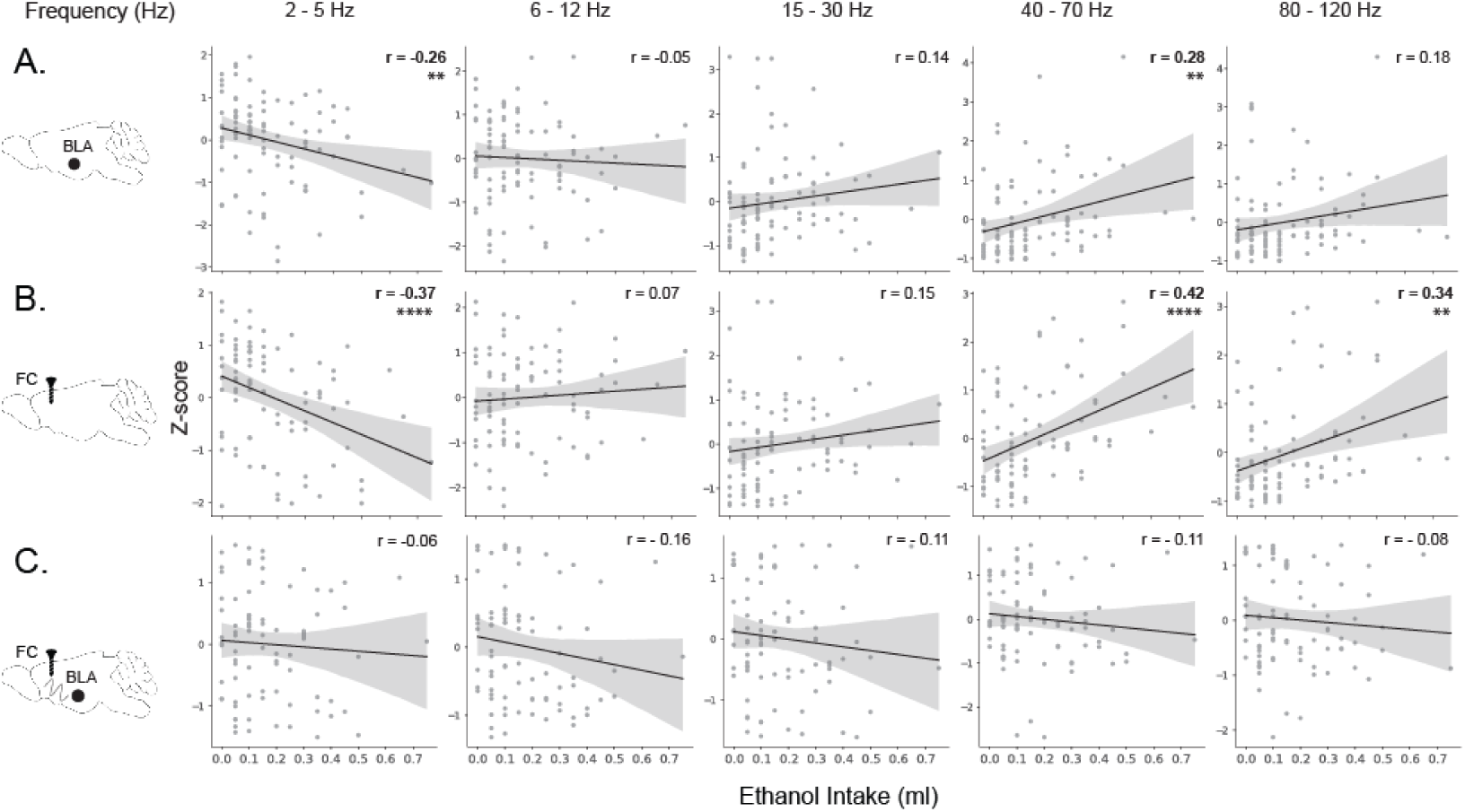
Power in BLA and frontal cortex but not coherence correlates with voluntary ethanol intake. Relationship between BLA power (n = 95 sessions (11 mice)) (A), frontal cortex power (n = 89 sessions (11 mice)) (B), coherence (z-score) between the BLA and frontal cortex (n = 84 sessions (11 mice)) (C), with the amount of ethanol consumed in the DID-MSA binge periods. Statistical analysis by Pearson correlation with permutation (see *Statistical Analysis* section). \*\**p* < 0.01, \*\*\*\**p* < 0.0001.

